# *Klebsiella quasipneumonaiae* provides a window into carbapenemase gene transfer, plasmid rearrangements and nosocomial acquisition from the hospital environment

**DOI:** 10.1101/486753

**Authors:** Amy J Mathers, Derrick Crook, Alison Vaughan, Katie Barry, Kasi Vegesana, Nicole Stoesser, Hardik Parikh, Robert Sebra, Shireen Kotay, A Sarah Walker, Anna E Sheppard

## Abstract

Several emerging pathogens have arisen as a result of selection pressures exerted by modern healthcare. *Klebsiella quasipneumoniae* was recently defined as a new species, yet its prevalence, niche, and propensity to acquire antimicrobial resistance genes are not fully described. We have been tracking inter- and intra-species transmission of the *Klebsiella pneumoniae* carbapenemase (KPC) gene, *bla*_KPC_, between bacteria isolated from a single institution. We applied a combination of Illumina and PacBio whole-genome sequencing to identify and compare *K. quasipneumoniae* from patients and the hospital environment over 10 and five-year periods respectively. There were 32 *bla*_KPC_-positive *K. quasipneumoniae* isolates, all of which were identified as *K. pneumoniae* in the clinical microbiology laboratory, from eight patients and 11 sink drains, with evidence for seven separate *bla*_KPC_ plasmid acquisitions. Analysis of a single subclade of *K. quasipneumoniae* subspecies *quasipneumoniae* (n=23 isolates) from three patients and six rooms demonstrated seeding of a sink by a patient, subsequent persistence of the strain in the hospital environment, and then probable transmission to another patient. Longitudinal analysis of this strain demonstrated the acquisition of two unique *bla*_KPC_ plasmids and then subsequent within-strain genetic rearrangement through transposition and homologous recombination. Our analysis highlights the apparent molecular propensity of *K. quasipneumoniae* to persist in the environment as well as acquire carbapenemase plasmids from other species and enabled an assessment of the genetic rearrangements which may facilitate horizontal transmission of carbapenemases.

## Introduction

In the last 50 years transformations in healthcare have created new niches for microorganisms such as *Acinetobacter baumannii* complex and *Candida auris* to arise from obscurity and emerge as important pathogens. Similarly, we have seen an increasing number of highly resistant *Klebsiella pneumoniae* strains which have been successfully transmitted worldwide(1). *Klebsiella pneumoniae* has proven to be an important contributor to the modern antibiotic resistance epidemic with its ability to acquire and carry antimicrobial resistance plasmids, as well as being successful a human pathogen. More recently, whole-genome sequencing has revealed that many isolates classified as *K. pneumoniae* actually encompass three related but distinct species – *K. pneumoniae*, *K. variicola* and *K. quasipneumoniae*(1, 2). *K. quasipneumoniae* was originally thought to be largely confined to agriculture and the environment, however it appears that it may also be prominent in human disease(3), and several recent reports have demonstrated that it harbors virulence factors and acquires clinically relevant genes of antimicrobial resistance(4, 5). Although there have been relatively few reports of *K. quasipneumoniae* to date, the true prevalence of this organism is likely underestimated as it is not generally distinguished from *K. pneumoniae* in routine testing of clinical laboratories(2).

Bacterial evolution via horizontal gene transfer is central to the ongoing crisis of antimicrobial resistance among clinically relevant bacteria. Hospital wastewater is being increasingly recognized as an ideal reservoir for resistance gene exchange and amplification, with ongoing antimicrobial selection pressure exerted through antimicrobials excreted in patient waste(6). Premise plumbing can be seeded by antimicrobial resistance genes in diverse bacterial strains and species, and represents a difficult-to-treat reservoir for ongoing gene exchange, creating successful drug-resistant bacteria that can thrive in both the environmental and human niches(7).

Whole-genome sequencing studies have demonstrated that our understanding of the interplay between antimicrobial resistance plasmids and their host strains/species is limited(8). The host range of a plasmid is critical for acquisition and persistence in specific species, but it appears that some bacterial strains are better equipped than others to prevent acquisition of or destroy foreign plasmid DNA(9). The durability of plasmid acquisition events and the creation of new highly resistant strains reflects complex dynamics which depend on the characteristics of the plasmid in question as well as host strain tolerance(10, 11). Seldom do we have the opportunity to witness strains acquiring plasmids *in vivo* or in the environment and inferences about genetic re-arrangements are often highly speculative. However, understanding the mechanisms and frequency of resistance gene transfer events occurring in real world contexts can provide important insights into the wider evolutionary landscape creating modern multidrug resistant bacteria which cannot be effectively modeled in lab experiments(12).

Within our institution we have seen ongoing transmission of diverse carbapenemase-producing organisms for the last decade, driven by genetic exchange of the *Klebsiella pneumoniae* carbapenemase (KPC) gene (*bla*_KPC_) in patients and the environment(13, 14). This has enabled us to understand specific pathways of genetic mobility involving numerous different mobile genetic elements and host bacterial species(13, 15). Herein we examine *bla*_KPC_ acquisition and associated genetic rearrangements within *K. quasipneumoniae* as a real-life representation of an emerging pathogen associated with the hospital wastewater environment.

## Results

From our collection of *bla*_KPC_-positive isolates from patients (2007-2017) and the hospital environment (2013-2017), there were a total of 32 *bla*_KPC_-positive *K. quasipneumoniae* isolates, all of which were identified as *K. pneumoniae* in the clinical microbiology laboratory (Table 1). Twenty-three of these were *K. quasipneumoniae* subspecies *quasipneumoniae* (KpIIA) (ten patient isolates from four patients and 13 environmental isolates from seven rooms) and nine were *K. quasipneumoniae* subspecies *similipneumoniae* (KpIIB) (five patient isolates from four patients and four environmental isolates from four rooms). The KpIIA and KpIIB isolates were separated by >100,000 single nucleotide variants (SNVs). We identified a single strain of KpIIA and four strains of KpIIB differing from each other by >20,000 SNVs (Fig. 1).

**Table 1.** All Sequenced *bla*_KPC_-*Klebsiella quasipneumoniae* isolates from patients and the hospital environment

| label | Isolate | Subspecies of *K. quasipneumoniae* | Date | Source | Infection/outcome |
| --- | --- | --- | --- | --- | --- |
| 1 | CAV1360 | *KpIIA* | Nov-09 | Sputum | Ventilator associated pneumonia. in complicated heart transplant recipient/Expired |
| 2 | CAV2013 | *KpIIA* | Nov-13 | Perirectal surveillance | N/A |
| 2 | CAVp203 | *KpIIA* | Dec-13 | Bronchoscopy | Ventilator associated pneumonia //Successful treatment |
| 2 | CAVp26 | *KpIIA* | Apr-14 | Blood | Intraabdominal infection/Expired |
| 2 | CAVp20 | *KpIIA* | Mar-14 | Perirectal surveillance | N/A |
| 2 | CAVp11 | *KpIIA* | Feb-14 | Abdominal fluid | Intraabdominal infection//Successful treatment |
| 2 | CAVp64 | *KpIIA* | Aug-14 | Perirectal surveillance | N/A |
| 2 | CAVp72 | *KpIIA* | Sep-14 | Perirectal surveillance | N/A |
| 2 | CAVp103 | *KpIIA* | Nov-14 | Blood | Successful treatment of ... |
| 3 | CAVp50 | *KpIIA* | Jul-14 | Perirectal surveillance | N/A |
| 3 | CAVp57 | *KpIIA* | Jul-14 | Perirectal surveillance | N/A |
| 3 | CAVp67 | *KpIIA* | Aug-14 | Perirectal surveillance | N/A |
| 3 | CAVp71 | *KpIIA* | Aug-14 | Perirectal surveillance | N/A |
| 3 | CAVp104 | *KpIIA* | Dec-14 | Perirectal surveillance | N/A |
| 4 | CAVp275 | *KpIIA* | Jul-15 | Urine | Complicated urinary tract infection/ Successful treatment |
| 5 | CAV1142 | *KpIIB* | Aug-09 | Perirectal surveillance | N/A |
| 6 | CAVp186 | *KpIIB* | Dec-13 | Perirectal surveillance | N/A |
| 7 | CAV2009 | *KpIIB* | Feb-14 | Perirectal surveillance | N/A |
| 8 | CAVp296 | *KpIIB* | Oct-15 | Perirectal surveillance | N/A |
| 8 | CAVp360 | *KpIIB* | Dec-16 | Perirectal surveillance | N/A |
| **Room A (MICU)** | CAV2244 | *KpIIA* | Jan-14 | Shower |  |
| **Room B (CTA)** | CAV2279 | *KpIIA* | Jan-14 | Shower |  |
| **Room C (STBICU)** | CAV1945 | *KpIIA* | Feb-14 | Drain swab (First sample after replacement) |  |
| **Room C (STBICU)** | CAV1947 | *KpIIA* | Feb-14 | p-trap water -(First sample after replacement) |  |
| **Room C (STBICU)** | CAV1964 | *KpIIA* | Mar-14 | Drain swab |  |
| **Room C (STBICU)** | CAV2018 | *KpIIA* | Apr-14 | p-trap water |  |
| **Room D (STBICU)** | CAV2019 | *KpIIA* | Apr-14 | p-trap water |  |
| **Room C (STBICU)** | CAV2397 | *KpIIA* | May-14 | Drain swab |  |
| **Room E (STBICU)** | CAV2697 | *KpIIA* | Jul-14 | Drain swab |  |
| **Room F (MICU)** | CAV2957 | *KpIIA* | Sep-15 | Drain swab |  |
| **Room G (STBICU)** | CAV2983 | *KpIIA* | Oct-15 | p-trap water |  |
| **Room G (STBICU)** | CAV2984 | *KpIIA* | Oct-15 | Drain swab |  |
| **Room G (STBICU)** | CAV3444 | *KpIIA* | Feb-16 | p-trap water |  |
| **Room H (MICU)** | CAV1880 | *KpIIB* | Dec-13 | Drain swab |  |
| **Room I (MICU)** | CAV1895 | *KpIIB* | Dec-13 | Drain swab |  |
| **Room J (STBICU)** | CAV1832 | *KpIIB* | Dec-13 | p-trap water |  |
| **Room K (STBICU)** | CAV1887 | *KpIIB* | Dec-13 | p-trap water |  |

**Fig. 1.**
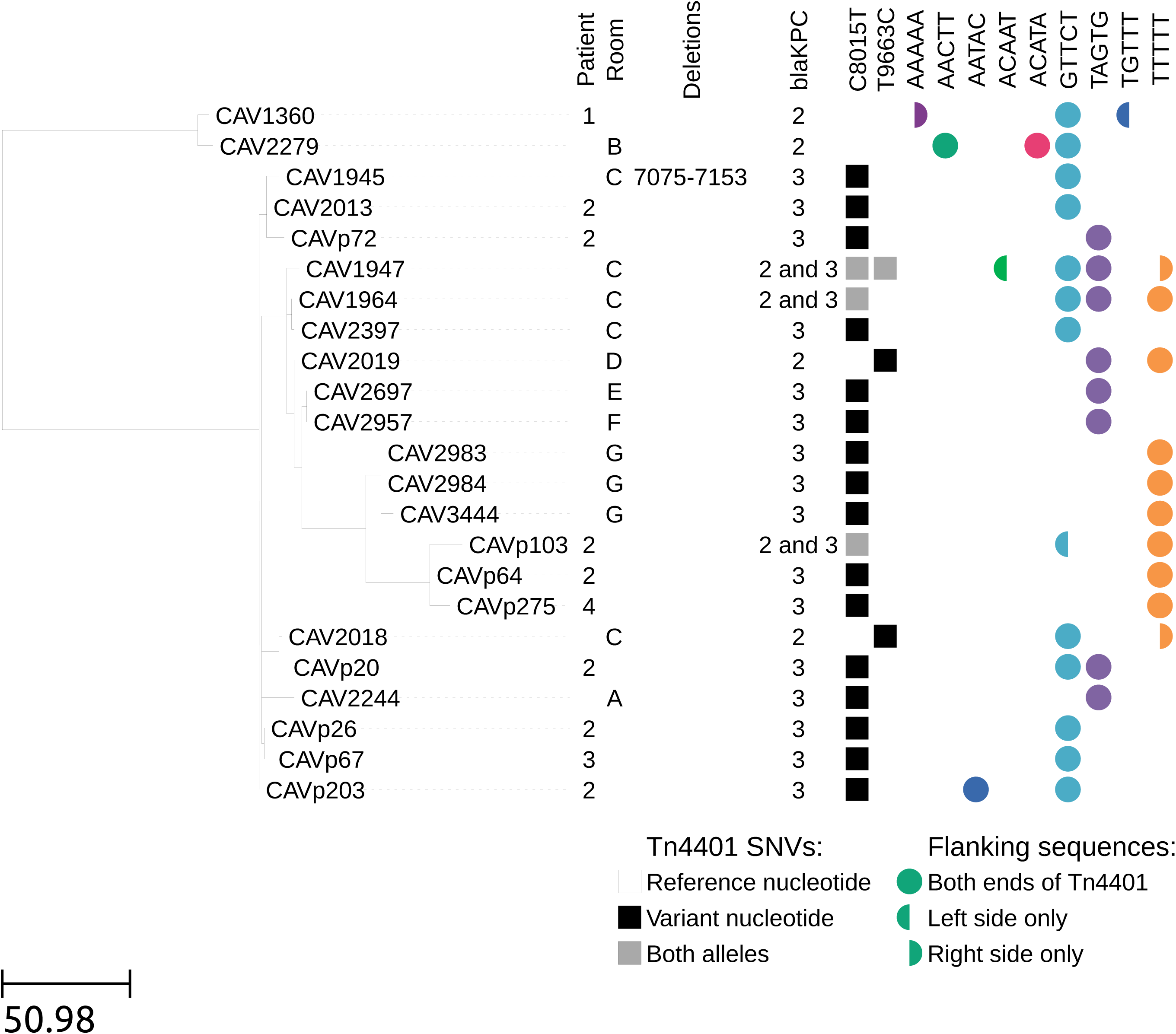
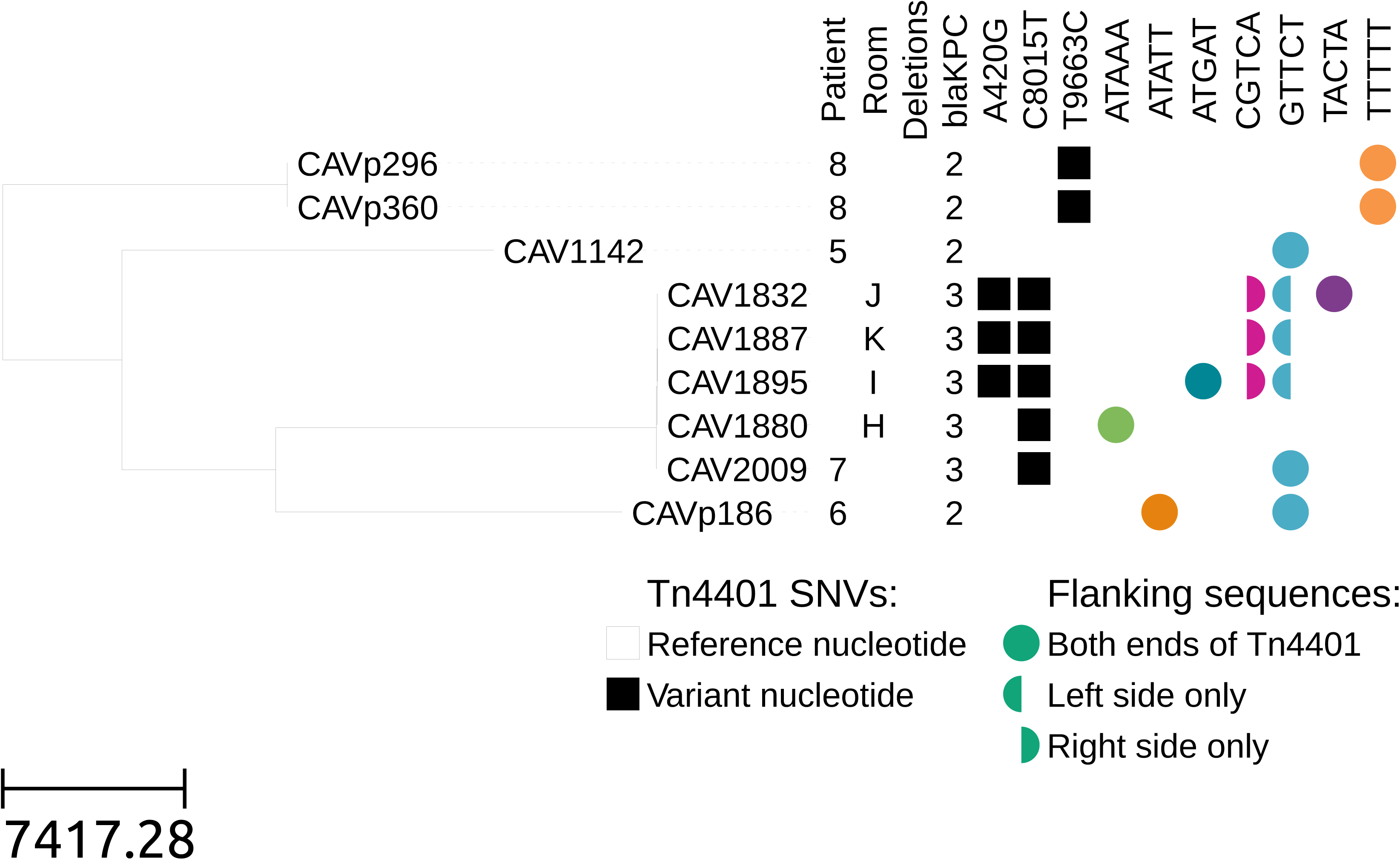
Maximum likelihood phylogeny for KpIIA (a) and KpIIB (b) isolates, with Tn*4401* variation and flanking genetic contexts. Branch lengths are shown as SNVs per genome.

Within the KpIIA strain, there were two subclades separated by ~150 SNVs (Fig. 1a). The first subclade contained two isolates separated by 10 SNVs (Fig. 1a). CAV1360 was from patient 1 in November 2009 and CAV2279 was identified in early 2014 (shortly after environmental sampling began) from room B that patient 1 had occupied in May 2009.

The second subclade of KpIIA contained isolates from three patients (patients 2-4) and six rooms (rooms A, C-G). The earliest of these was from patient 2 in November 2013. Patient 2 was in the hospital with a prolonged stay in the Surgical Trauma and Burn Intensive Care Unit (STBICU) following complications of a liver transplant (Figure 2). Patient 2 was noted to be first colonized with *bla*_KPC_-positive KpIIA in November 2013. KpIIA was not found in the STBICU environment prior to closure for remediation of KPC-contamination of the drains in December 2013. Following drain exchange and unit re-opening patient 2 was immediately moved back into the STBICU and subsequently occupied rooms C, D, E and G in the STBICU, suggesting that the KpIIA isolates in these rooms originated from patient 2 (Figure 2). Patient 3 was admitted to the STBICU at the same time as patient 2 and thus could have acquired KPC-KpIIA directly from patient 2 without environmental transmission. Patient 4 was later admitted to STBICU room E for 28 days and discharged before he was found to have KpIIA. He was never on a ward at the same time as any other patients known to carry KpIIA, suggesting acquisition from the hospital environment.

**Fig. 2.**
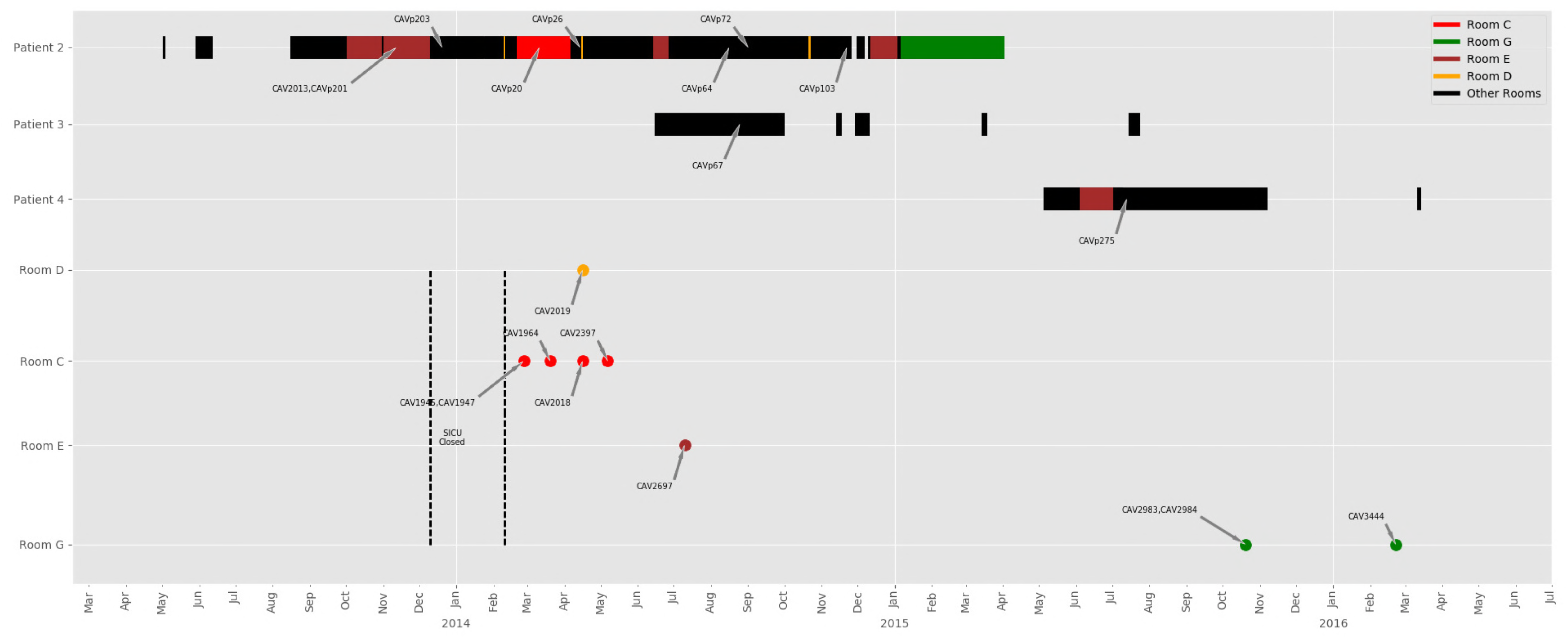
Patient movements and positive environmental samples with a single strain of *K. quasipneumoniae* (KpIIA) in the STBICU. Colored bars for patients match rooms where environmental isolates were identified. Black bars represent rooms with no KpIIA identified. The dotted lines indicate STBICU closure with removal and new installation of sink drains and exposed sink plumbing. Patient 1 is not depicted as there was no admission to the STBICU and no overlap in time or space with other patients carrying KpIIA.

There were four patients (patients 5-8) carrying four distinct strains of *bla*_KPC_-KpIIB seen over a five year period (Fig. 1b, Table 1). For patient 7, the same KpIIB strain (~80 SNV differences) was also seen in sinks from two rooms in the Medical Intensive Care Unit (MICU) (rooms H-I) and two rooms in the STBICU (rooms J-K) in December 2013 when environmental sampling first began; this preceded detection in the patient in February 2014. Patient 7 was admitted to the MICU, but did not stay in rooms H or I. The other three patients with KpIIB each had a unique *bla*_KPC_ strain, none of which were identified in another patient or the environment. Patient 6 had a prolonged hospital stay and was also colonized/infected with another *bla*_KPC_-positive species (*K. pneumoniae*).

Three patients developed infections with KPC-KpIIA (Table 1). Patient 1 died of ventilator-associated pneumonia with KPC-KpIIA following a complicated heart transplant. Patient 2 had both ventilator-acquired pneumonia, which was successfully treated, and a subsequent untreatable intraabdominal infection with KPC-KpIIA bacteremia, which contributed to the patient’s death after a long hospital stay with a complicated liver transplant. Patient 4 had a successfully treated complicated KPC-KpIIA urinary tract infection. Patient 3 did not develop an infection with KpIIA. None of the patients with KpIIB developed *K. quasipneumoniae* infections, however two of the patients did develop infections with other species carrying *bla*_KPC_ (*K. pneumoniae* for patient 6 and *Serratia marcescens* for patient 8) (Table 2).

**Table 2.**
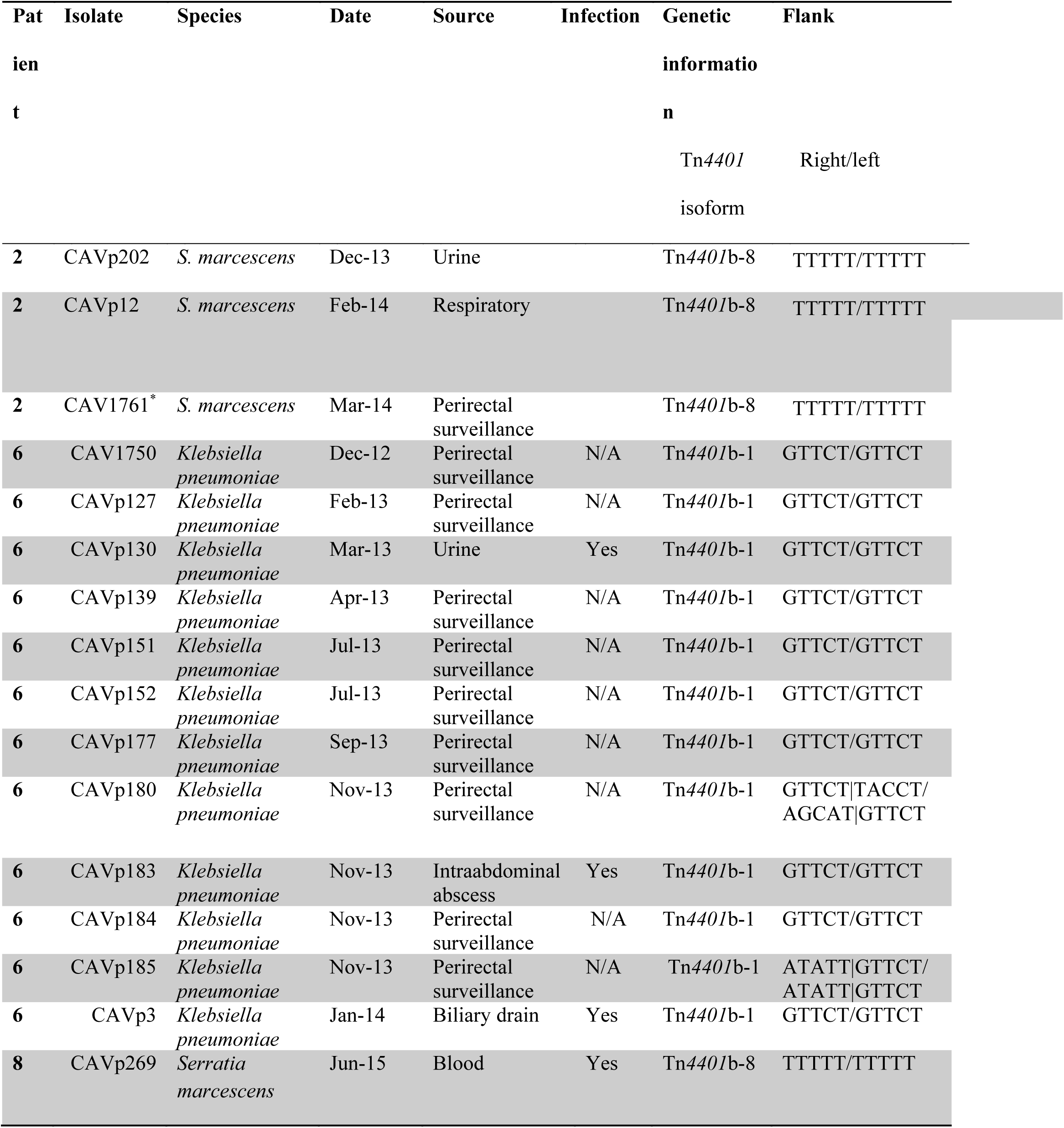
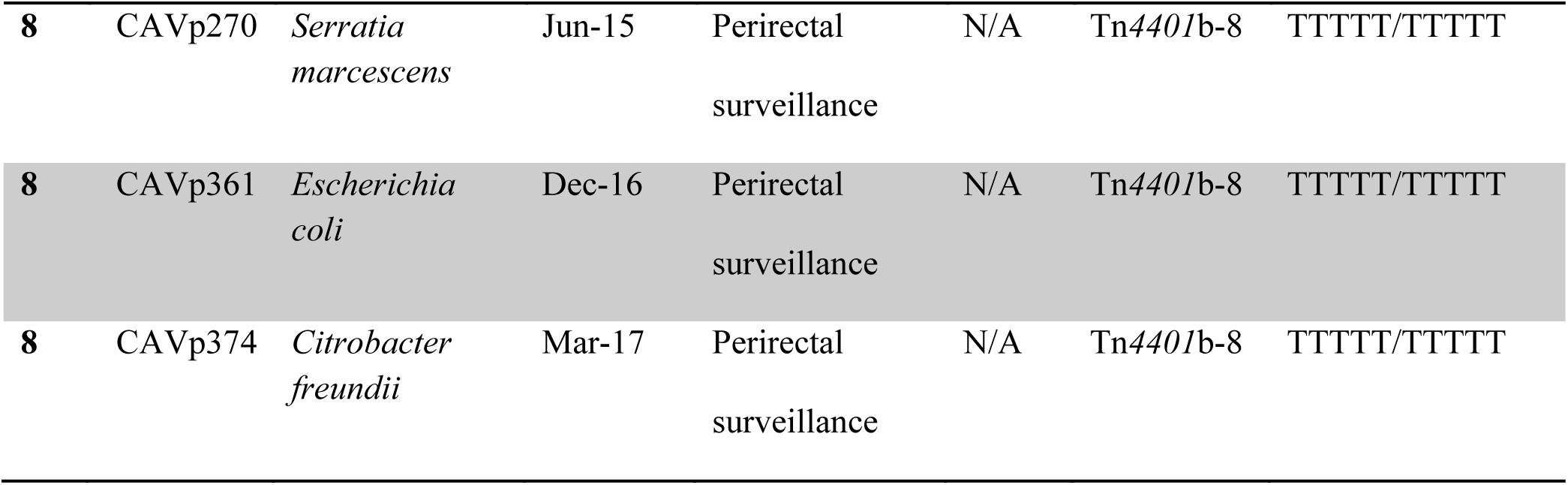
All additional *bla*_KPC_-positive isolates from patients with *K. quasipneumoniae*

### Genetic variation and rearrangements within KpIIA

All KpIIA isolates were closely related at the core chromosome level, with a maximum divergence of <180 SNVs. If *bla*_KPC_ were acquired only once in this lineage then any sequence variation within the 10 kb *bla*_KPC_ transposon Tn*4401* would be the result of mutational change, which is expected to be rare. Surprisingly, the Illumina sequence data revealed a great deal of sequence variation within Tn*4401* (Fig. 1a). Two sites (positions 8015 and 9663 in the Tn*4401*b reference) showed variation at the single-nucleotide level, and one isolate had a deletion at positions 7075-7153. Interestingly, several isolates showed mixtures at one or both of the variable sites, indicating two or more different versions of Tn*4401* in the same isolate. This included mixtures at position 8015, which is located within the *bla*_KPC_ gene and differentiates *bla*_KPC-2_ and *bla*_KPC-3_, indicating that there were isolates with both *bla*_KPC_ alleles.

Similarly, if a single *bla*_KPC_ plasmid were acquired and stably maintained within KpIIA, then we would expect to see a single flanking sequence context for Tn*4401*. On the contrary, there was significant diversity in Tn*4401* flanking regions, with eight and seven different 5 bp sequences on the left and right sides of Tn*4401* respectively, suggesting active transposition of Tn*4401* within KpIIA and/or multiple plasmid acquisitions.

To better understand the origin of the genetic diversity within and surrounding Tn*4401*, we performed long-read PacBio sequencing on three of the KpIIA isolates (CAV2013 from patient 2, CAV1947 from room C and CAV2018 from room C), as well as a *S. marcescens* isolate from patient 2 (CAV1761). The room C isolates were chosen because this room only became positive after admission of patient 2 following sink trap exchange in the STBICU, hence they are expected to be descended from the patient 2 KpIIA.

Both patient 2 isolates had a single *bla*_KPC_ plasmid each (Figure 3a-b). The KpIIA isolate had a 47,095 bp “RepA” *bla*_KPC-3_ plasmid, and the *S. marcescens* isolate had a 69,158 bp IncU/IncX5 *bla*_KPC-2_ plasmid (16). Both plasmids contained Tn*4401*b, however there were two SNV differences within the Tn*4401*b sequence, one at position 8015 (differentiating *bla*_KPC-2_ and *bla*_KPC-3_) and one at position 9663.

**Fig. 3.**
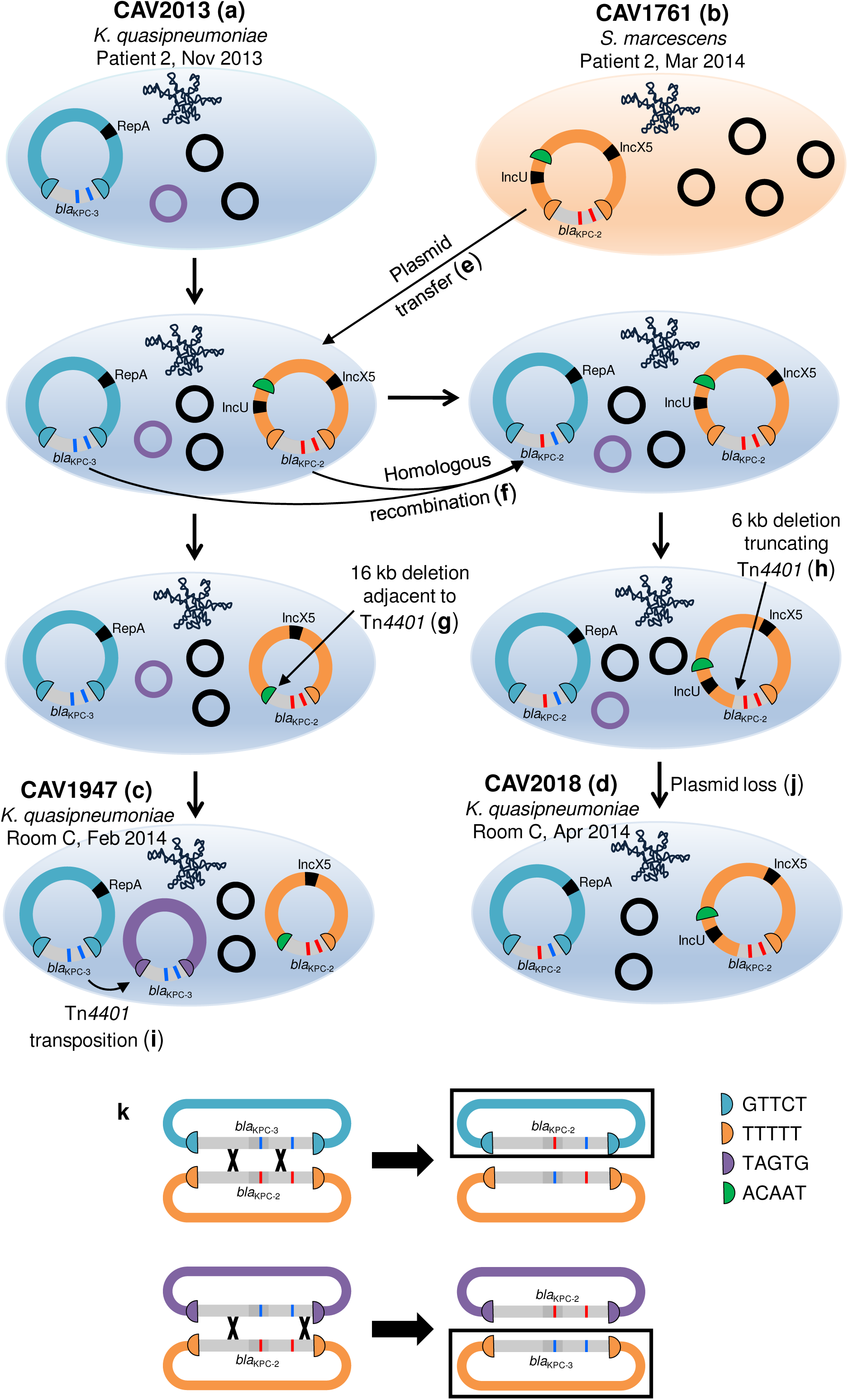
Plasmid structures determined from long-read sequencing of four isolates and inferred intermediate *bla*_KPC_ plasmid structures. a-d. Sequenced isolates. e-j. Inferred intermediate plasmid structures. Note that the ordering of deletion, homologous recombination, transposition and plasmid loss events is arbitrarily represented as the actual order of events is unknown. k. Examples of crossover events leading to the generation of new combinations of SNVs within Tn*4401* (top) or the complete swapping of Tn*4401* variants between different plasmids (bottom). Black boxes indicate products of homologous recombination that were observed in long-read data (top) or Illumina data (bottom).

The KpIIA isolates from room C (CAV1947 and CAV2018) had three and two *bla*_KPC_ plasmids respectively (Fig. 3c-d). Both isolates harbored the IncU/IncX5 *bla*_KPC_ plasmid from the patient 2 *S. marcescens* isolate, indicating likely *bla*_KPC_ plasmid transfer from *S. marcescens* to *K. quasipneumoniae* (Fig. 3e). In CAV1947, the plasmid sequence was identical to the patient isolate, CAV1761, with the exception of two large indels (Fig. 4a). One of these was a 16,315 bp deletion immediately adjacent to Tn*4401*, presumably as a result of intramolecular transposition, that converted the left flanking sequence from TTTTT to ACAAT and removed the IncU replicon sequence (Fig. 3g). In CAV2018, the plasmid sequence was identical to CAV1761, except for a single 5,923 bp deletion that truncated part of the Tn*4401* sequence (Fig. 3h, 4a).

**Fig 4.**
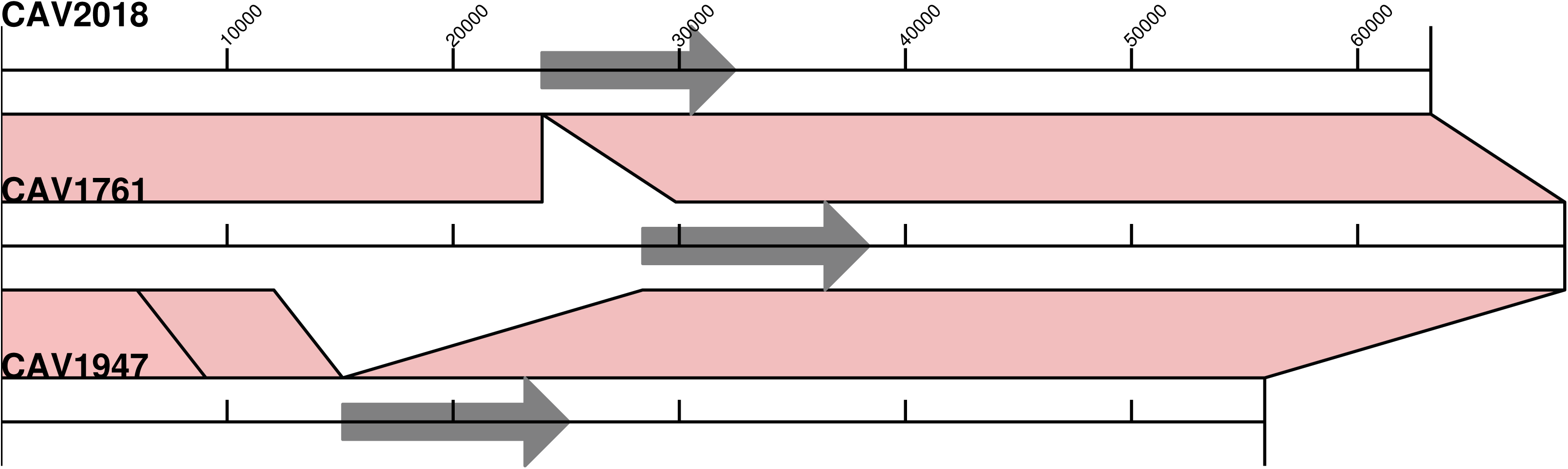
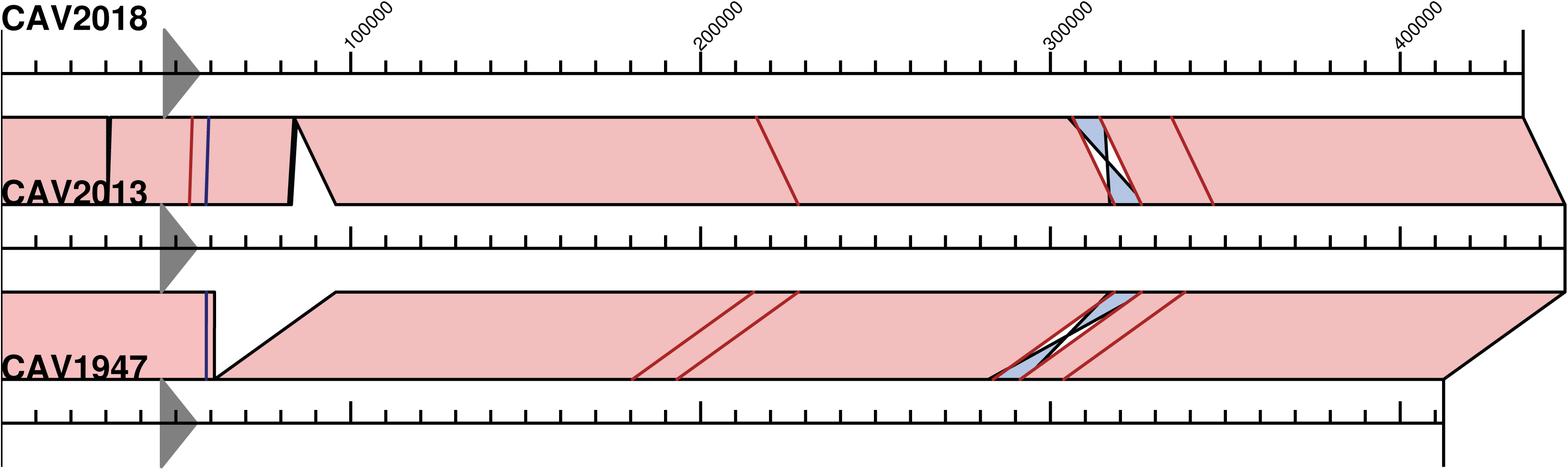
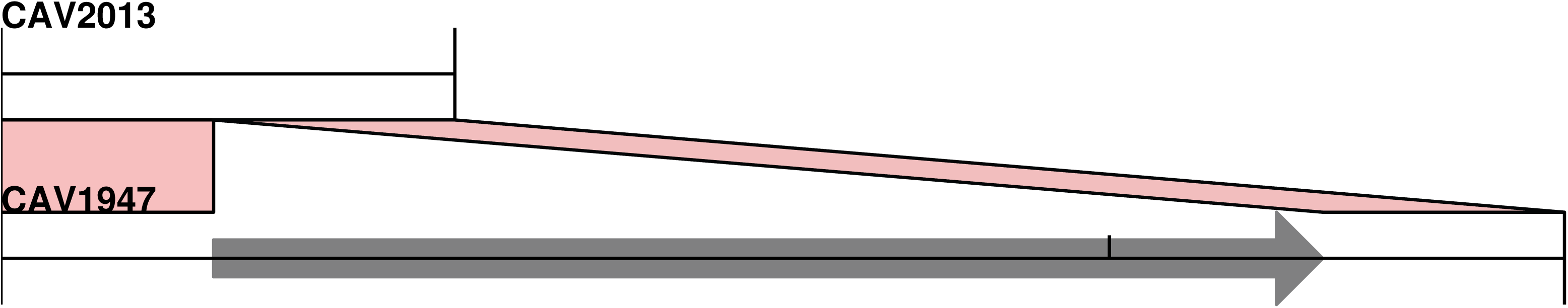
Alignments of IncU/IncX5 (a), RepA (b) and non-typeable (c) *bla*_KPC_ plasmid structures determined from long-read sequencing. Tn*4401* is indicated by a grey arrow. Light pink shading indicates regions of identity, light blue shading shows inverted regions, SNVs are indicated by red lines and short indels by blue lines.

Both isolates also harboured the ancestral RepA *bla*_KPC_ plasmid from the patient 2 KpIIA isolate, with several SNVs and large indels (Fig. 4b). Interestingly, in CAV2018, one of the SNVs was located within Tn*4401*, such that the CAV2018 RepA plasmid contained *bla*_KPC-2_ rather than *bla*_KPC-3_. Given that there was plasmid transfer of the IncU/IncX5 *bla*_KPC-2_ plasmid from *S. marcescens*, we infer that the *bla*_KPC-2_-containing RepA plasmid most likely arose as a result of homologous recombination between these two different plasmids flanking the *bla*_KPC_ region (Fig. 3f, k). The Illumina data also revealed similar patterns of homologous recombination in other isolates (notably CAV2983, CAV2984, CAV3444, CAVp64 and CAVp275, which all have the TTTTT IncU/IncX5 plasmid flanking sequences, but with the C8015T *bla*_KPC-3_ mutation and without the T9663C mutation), suggesting frequent exchange of Tn*4401* variants between different *bla*_KPC_ plasmids within the same host bacterium (Fig. 1, 3k).

CAV1947 also harboured a third *bla*_KPC_ plasmid, representing transposition of Tn*4401* into a 4,095 bp non-typeable plasmid that was present in the CAV2013 ancestor from patient 2 (Fig. 3i, 4c).

### K. quasipneumoniae has acquired *bla*_KPC_ on multiple occasions

Within KpIIB, there were four divergent strains separated by >20,000 SNVs, suggesting four separate acquisitions of *bla*_KPC_ in this subspecies. Within KpIIA, there were two subclades separated by ~180 SNVs. Given that Tn*4401* variation and flanking sequences were different between the two subclades (apart from the GTTCT flanking sequence which is known to be present in many different *bla*_KPC_ plasmids)(13); and that there was no epidemiological overlap, it is most likely that the subclades acquired *bla*_KPC_ independently. Additionally, as described above, the second subclade likely acquired *bla*_KPC_ on two occasions, with the second acquisition originating from *S. marcescens*. Therefore, overall there were likely seven acquisitions of *bla*_KPC_ by *K. quasipneumoniae*, three in KpIIA and four in KpIIB.

Interestingly, there was evidence that one of the acquisitions in KpIIB also originated from *S. marcescens*, indicating the compatibility of these two species in exchanging plasmids. This was in the patient 8 KpIIB lineage. Patient 8 was first colonised with *bla*_KPC_-*S. marcescens* carrying Tn*4401*b with a T9663C mutation and TTTTT/TTTTT flanking sequences. Four months later, *bla*_KPC_-KpIIB was identified with the same Tn*4401* mutation and flanking sequences, suggesting plasmid transfer from *S. marcescens* to *K. quasipneumoniae* within this patient.

## Discussion

We describe the behaviour of nosocomial *bla*_KPC_-positive *K. quasipneumoniae* strains within a single-hospital setting, observing their propensity to uptake multiple carbapenemase plasmids from other species, and disseminate between patients and sink drains. Our study also suggests that rapid genetic rearrangement occurs in the mobile genetic elements carrying *bla*_KPC_ in KpIIA.

There is increasing recognition that the hospital environment is an important potential reservoir in the transmission of carbapenemase-producing *Enterobacteriaceae* (CPE), but delineating transmission chains is often challenging(17, 18). Through our *K. quasipneumoniae* example we provide compelling evidence for patient-to-drain and drain-to-patient transmission, as has been observed in other studies(7). We also provide evidence supporting the ability of *K. quasipneumoniae* to be maintained in the environment for a long period of time, with the first subclade of KpIIA detected in the environment on initial sampling, even though it had not been seen in a patient nor had that patient been in the room for over three years. The costly closure of the STBICU and exchange of all the sink drain plumbing pipes had a limited effect on environmental contamination with CPE; instead it appears to have provided an environment for immediate new seeding and establishment of previously unobserved carbapenem-resistant strains. Understanding the dynamics and natural history of colonization of premise plumbing with CPE will be important in designing effective interventions to limit transmission(19).

Although there have only been a handful of reports of *K. quasipneumoniae* since its definition as a species in 2014, it does appear that this organism is widespread(2, 5, 20, 21). As seen here, it is not readily distinguished from *K. pneumoniae* with current clinical microbiology techniques and thus the true prevalence is unknown(2, 22). On the evolutionary time scale, modern medicine has provided a novel ecology with immunocompromised patients, widespread antimicrobial use, newly circulating antimicrobial resistance genes and the design of the modern hospital providing new microbiologic niche for organisms to emerge(7, 23). As seen here we provide evidence for *K. quasipneumoniae* to be sustained in both a human host and the environment encountering several different species which may be relatively new in the evolutionary tree of *Klebsiella* sp. As a consequence of these encounters transfer of mobile DNA occurs via traceable carbapenemase plasmids. We found evidence for seven independent acquisitions of *bla*_KPC_ by *K. quasipneumoniae*, suggesting that this species is amenable to plasmid uptake from other species of Enterobacteriaceae. Given the difficulties in accurately identifying *K. quasipneumoniae*, this species may therefore be more significant in the context of *bla*_KPC_ dissemination than has previously been recognised.

Within *K. quasipneumoniae*, there was surprising variability in mobile elements carrying *bla*_KPC_, which was the result of several different processes observed amongst a limited number of highly related isolates(n=23). Specifically, there were multiple independent *bla*_KPC_ plasmid acquisitions, homologous recombination between different *bla*_KPC_ plasmids, transposition of Tn*4401* into new plasmids, intramolecular transposition of Tn*4401*, a deletion within Tn*4401* and a deletion truncating Tn*4401*. This high degree of genetic mobility has been similarly observed in other small studies(25, 26), and highlights the difficulty in developing an accurate understanding of the transmission epidemiology of important drug resistance genes which can be rapidly mobilized by multiple independent genetic modalities.

Within KpIIA, there were multiple acquisitions of *bla*_KPC_ within the same lineage, such that a *bla*_KPC_-positive KpIIA strain acquired a second, unrelated *bla*_KPC_ plasmid from *S. marcescens*. Consequently, there were then two different *bla*_KPC_ plasmids, with different Tn*4401* sequences and different *bla*_KPC_ alleles, within the same host bacterium. This situation facilitated multiple rearrangements via homologous recombination between the different plasmids, resulting in the generation of new combinations of Tn*4401* SNVs and host plasmids. Multiple acquisition of resistance plasmids followed by rearrangements between those plasmids is likely to be important in the generation of adaptive allelic combinations which contribute to the amplification of cross-class antimicrobial resistance within strains. High-risk clones with a propensity for uptake of antimicrobial resistance plasmids may represent important targets for intervention(24).

This study has several limitations. Most notably, it is a small retrospective series, preventing a full understanding of the role of the environment. Also, the order of genetic rearrangements is also not completely known given the limited number of long-read sequenced isolates and inability to capture all isolates from the environment over time. We offer however, that this is higher resolution than seen in many studies, and the analysis does contribute to the greater understanding of rapid rearrangement and mechanisms at play around mobility of genetic elements harbouring genes of antibiotic resistance in *Enterobacteriaceae*.

In summary, we demonstrate the relevance of *K. quasipneumoniae* as a species fit for nosocomial transmission in the modern era that is capable of acquiring and maintaining relevant resistance elements.

## Methods

### Setting

Isolates were collected at the University of Virginia, a 619-bed tertiary care hospital, from August 2007-May 2017. A robust *K. pneumoniae* carbapenemase-producing organism (KPCO) prevention program existed throughout the study period as previously described(27), and included perirectal screening beginning in April 2009 in the medical intensive care unit (MICU) and surgical intensive care unit (STBICU), and weekly screening of all patients in the MICU and STBICU as well as units where any known KPCO-colonized patient was present(28). Screening was performed as previously described(28). Clinical Enterobacteriales and Aeromonadaceae isolates, as identified by MALDI-TOF or VITEK2 (Biomerieux, Durham, NC), with an elevated ertapenem or meropenem minimum inhibitory concentration (MIC) by VITEK2 (Biomerieux, Durham, NC) immediately underwent CarbaR (Cepheid Sunnyvale, CA) carbapenemase PCR testing. All species identification was performed using a combination of VITEK2, VITEK-MS (Biomerieux, Durham, NC). Clinical data was gathered by retrospective electronic medical record review under University of Virginia Health Sciences IRB#13558 with waiver of consent.

In September 2013 sink trap sampling for KPCO began using previously described techniques(14) with a swab for drain collection and p-trap water. Following identification of KPCO in the hospital environment, the STBICU was closed to patient care in December 2013. Over the following 9 weeks all sink drain pipes were removed and replaced with sink traps that eliminated overflows on the sink bowl. Patients were readmitted to the surgical intensive care unit in February 2014. Bleach, hydrogen peroxide and ozone impregnated water (2ppm) were applied weekly from February-May 2014 in the STBICU (following drain exchange and sink bowl overflow closure and removal) and from March-May 2014 in the MICU (without drain exchange or sink bowl overflow removal).

### Whole-genome sequencing

Illumina sequencing was performed as described previously(29). PacBio long-read sequencing and assembly were performed as previously described(13).

Broad level species classification was performed using Kraken(30). To identify *K. quasipneumoniae* isolates, we queried all isolates initially classified as *K. pneumoniae* against reference sequences representing each of the four clades in Holt et al(1). We arbitrarily selected a single reference sequence for each clade; these were: ERR025521 (KpI), ERR025986 (KpIIA), ERR025528 (KpIIB) and ERR025573 (KpIII). We used mash v1.1.1(31) with parameters “-r -m 5” to compare Illumina data for each of our isolates to these reference sequences. Each isolate was then assigned to one of the four Kp clades according to the reference with the lowest distance value. All isolates assigned to KpIIA or KpIIB were included in the analysis. In addition, we also included any other KPCO isolates from patients carrying *K. quasipneumoniae*.

To identify chromosomal single-nucleotide variants (SNVs), Illumina reads for each *K. quasipneumoniae* isolate were mapped to the CAV2013 chromosome sequence (derived from long-read sequencing), with mapping and variant calling performed as described previously (32). A phylogeny was generated using IQ-TREE v1.3.13 (33) from an alignment of variable sites where at least 70% of samples had a high-quality reference/variant call (i.e. we excluded sites where >30% of samples had an “N” call). This was run with parameters “-blmin 0.000000001 -nt 4 -m GTR”, with -fconst used to specify the number of invariant sites.

To identify Tn*4401* variation and flanking sequences from Illumina data, we used TETyper with published parameters(34).

Plasmid Inc typing was performed using the February 2018 version of the PlasmidFinder Enterobacteriaceae database(16), with an identity threshold of 95% and minimum length 60%.

## Funding

This work was funded in part by a contract from the Centres for Disease Control and Prevention (CDC) Broad Agency Announcement BAA 200-2017-96194. ASW, DWC, ASW are affiliated to the National Institute for Health Research Health Protection Research Unit (NIHR HPRU) in Healthcare Associated Infections and Antimicrobial Resistance at University of Oxford in partnership with Public Health England (PHE) [grant HPRU-2012-10041] and are supported by the Oxford NIHR Biomedical Research Centre. The views expressed are those of the author(s) and not necessarily those of the NHS, the NIHR, the Department of Health or Public Health England.

## References

1. Holt KE, Wertheim H, Zadoks RN, Baker S, Whitehouse CA, Dance D, Jenney A, Connor TR, Hsu LY, Severin J, Brisse S, Cao H, Wilksch J, Gorrie C, Schultz MB, Edwards DJ, Nguyen KV, Nguyen TV, Dao TT, Mensink M, Minh VL, Nhu NT, Schultsz C, Kuntaman K, Newton PN, Moore CE, Strugnell RA, Thomson NR. 2015. Genomic analysis of diversity, population structure, virulence, and antimicrobial resistance in Klebsiella pneumoniae, an urgent threat to public health. Proc Natl Acad Sci U S A 112:E3574–81.

2. Long SW, Linson SE, Ojeda Saavedra M, Cantu C, Davis JJ, Brettin T, Olsen RJ. 2017. Whole-Genome Sequencing of Human Clinical Klebsiella pneumoniae Isolates Reveals Misidentification and Misunderstandings of Klebsiella pneumoniae, Klebsiella variicola, and Klebsiella quasipneumoniae. mSphere 2.

3. Brisse S, Passet V, Grimont PA. 2014. Description of Klebsiella quasipneumoniae sp. nov., isolated from human infections, with two subspecies, Klebsiella quasipneumoniae subsp. quasipneumoniae subsp. nov. and Klebsiella quasipneumoniae subsp. similipneumoniae subsp. nov., and demonstration that Klebsiella singaporensis is a junior heterotypic synonym of Klebsiella variicola. Int J Syst Evol Microbiol 64:3146–52.

4. Becker L, Fuchs S, Pfeifer Y, Semmler T, Eckmanns T, Korr G, Sissolak D, Friedrichs M, Zill E, Tung ML, Dohle C, Kaase M, Gatermann S, Russmann H, Steglich M, Haller S, Werner G. 2018. Whole Genome Sequence Analysis of CTX-M-15 Producing Klebsiella Isolates Allowed Dissecting a Polyclonal Outbreak Scenario. Front Microbiol 9:322.

5. Nicolas MF, Ramos PIP, Marques de Carvalho F, Camargo DRA, de Fatima Morais Alves C, Loss de Morais G, Almeida LGP, Souza RC, Ciapina LP, Vicente ACP, Coimbra RS, Ribeiro de Vasconcelos AT. 2018. Comparative Genomic Analysis of a Clinical Isolate of Klebsiella quasipneumoniae subsp. similipneumoniae, a KPC-2 and OKP-B-6 Beta-Lactamases Producer Harboring Two Drug-Resistance Plasmids from Southeast Brazil. Front Microbiol 9:220.

6. Hocquet D, Muller A, Bertrand X. 2016. What happens in hospitals does not stay in hospitals: antibiotic-resistant bacteria in hospital wastewater systems. J Hosp Infect 93:395–402.

7. Weingarten RA, Johnson RC, Conlan S, Ramsburg AM, Dekker JP, Lau AF, Khil P, Odom RT, Deming C, Park M, Thomas PJ, Henderson DK, Palmore TN, Segre JA, Frank KM. 2018. Genomic Analysis of Hospital Plumbing Reveals Diverse Reservoir of Bacterial Plasmids Conferring Carbapenem Resistance. MBio 9.

8. Suzuki H, Yano H, Brown CJ, Top EM. 2010. Predicting plasmid promiscuity based on genomic signature. J Bacteriol 192:6045–55.

9. Price VJ, Huo W, Sharifi A, Palmer KL. 2016. CRISPR-Cas and Restriction-Modification Act Additively against Conjugative Antibiotic Resistance Plasmid Transfer in Enterococcus faecalis. mSphere 1.

10. Loftie-Eaton W, Yano H, Burleigh S, Simmons RS, Hughes JM, Rogers LM, Hunter SS, Settles ML, Forney LJ, Ponciano JM, Top EM. 2016. Evolutionary Paths That Expand Plasmid Host-Range: Implications for Spread of Antibiotic Resistance. Mol Biol Evol 33:885–97.

11. Loftie-Eaton W, Bashford K, Quinn H, Dong K, Millstein J, Hunter S, Thomason MK, Merrikh H, Ponciano JM, Top EM. 2017. Compensatory mutations improve general permissiveness to antibiotic resistance plasmids. Nat Ecol Evol 1:1354–1363.

12. Hardiman CA, Weingarten RA, Conlan S, Khil P, Dekker JP, Mathers AJ, Sheppard AE, Segre JA, Frank KM. 2016. Horizontal Transfer of Carbapenemase-Encoding Plasmids and Comparison with Hospital Epidemiology Data. Antimicrob Agents Chemother 60:4910–9.

13. Sheppard AE, Stoesser N, Wilson DJ, Sebra R, Kasarskis A, Anson LW, Giess A, Pankhurst LJ, Vaughan A, Grim CJ, Cox HL, Yeh AJ, Sifri CD, Walker AS, Peto TE, Crook DW, Mathers AJ, Group MMMMI. 2016. Nested Russian Doll-like Genetic Mobility Drives Rapid Dissemination of the Carbapenem Resistance Gene blaKPC. Antimicrob Agents Chemother.

14. Mathers AJ, Vegesana K, German Mesner I, Barry KE, Pannone A, Baumann J, Crook DW, Stoesser N, Kotay S, Carroll J, Sifri CD. 2018. Intensive Care Unit Wastewater Interventions to Prevent Transmission of Multi-species Klebsiella pneumoniae Carbapenemase (KPC) Producing Organisms. Clin Infect Dis.

15. Mathers AJ, Stoesser N, Chai W, Carroll J, Barry K, Cherunvanky A, Sebra R, Kasarskis A, Peto TE, Walker AS, Sifri CD, Crook DW, Sheppard AE. 2017. Chromosomal Integration of the Klebsiella pneumoniae Carbapenemase Gene, blaKPC, in Klebsiella Species Is Elusive but Not Rare. Antimicrob Agents Chemother 61.

16. Carattoli A, Zankari E, Garcia-Fernandez A, Voldby Larsen M, Lund O, Villa L, Moller Aarestrup F, Hasman H. 2014. In silico detection and typing of plasmids using PlasmidFinder and plasmid multilocus sequence typing. Antimicrob Agents Chemother 58:3895–903.

17. Kotsanas D, Wijesooriya WR, Korman TM, Gillespie EE, Wright L, Snook K, Williams N, Bell JM, Li HY, Stuart RL. 2013. "Down the drain": carbapenem-resistant bacteria in intensive care unit patients and handwashing sinks. Med J Aust 198:267–9.

18. De Geyter D, Blommaert L, Verbraeken N, Sevenois M, Huyghens L, Martini H, Covens L, Piérard D, Wybo I. 2017. The sink as a potential source of transmission of carbapenemase-producing Enterobacteriaceae in the intensive care unit. Antimicrob Resist Infect Control 6:24.

19. Hopman J, Tostmann A, Wertheim H, Bos M, Kolwijck E, Akkermans R, Sturm P, Voss A, Pickkers P, Vd Hoeven H. 2017. Reduced rate of intensive care unit acquired gram-negative bacilli after removal of sinks and introduction of 'water-free' patient care. Antimicrob Resist Infect Control 6:59.

20. Gan HM, Rajasekaram G, Eng WWH, Kaniappan P, Dhanoa A. 2017. Whole-Genome Sequences of Two Carbapenem-Resistant Klebsiella quasipneumoniae Strains Isolated from a Tertiary Hospital in Johor, Malaysia. Genome Announc 5.

21. Shankar C, Nabarro LEB, Muthuirulandi Sethuvel DP, Raj A, Devanga Ragupathi NK, Doss GP, Veeraraghavan B. 2017. Draft genome of a hypervirulent Klebsiella quasipneumoniae subsp. similipneumoniae with novel sequence type ST2320 isolated from a chronic liver disease patient. J Glob Antimicrob Resist 9:30–31.

22. Martinez-Romero E, Rodriguez-Medina N, Beltran-Rojel M, Silva-Sanchez J, Barrios-Camacho H, Perez-Rueda E, Garza-Ramos U. 2018. Genome misclassification of Klebsiella variicola and Klebsiella quasipneumoniae isolated from plants, animals and humans. Salud Publica Mex 60:56–62.

23. Gomi R, Matsuda T, Yamamoto M, Chou PH, Tanaka M, Ichiyama S, Yoneda M, Matsumura Y. 2018. Characteristics of Carbapenemase-Producing Enterobacteriaceae in Wastewater Revealed by Genomic Analysis. Antimicrob Agents Chemother.

24. Mathers AJ, Peirano G, Pitout JD. 2015. The Role of Epidemic Resistance Plasmids and International High-Risk Clones in the Spread of Multidrug-Resistant Enterobacteriaceae. Clin Microbiol Rev 28:565–591.

25. Wailan AM, Sartor AL, Zowawi HM, Perry JD, Paterson DL, Sidjabat HE. 2015. Genetic Contexts of blaNDM-1 in Patients Carrying Multiple NDM-Producing Strains. Antimicrob Agents Chemother 59:7405–10.

26. Snesrud E, Ong AC, Corey B, Kwak YI, Clifford R, Gleeson T, Wood S, Whitman TJ, Lesho EP, Hinkle M, McGann P. 2017. Analysis of Serial Isolates of mcr-1-Positive Escherichia coli Reveals a Highly Active ISApl1 Transposon. Antimicrob Agents Chemother 61.

27. Grabowski ME, Kang H, Wells KM, Sifri CD, Mathers AJ, Lobo JM. 2017. Provider Role in Transmission of Carbapenem-Resistant Enterobacteriaceae. Infect Control Hosp Epidemiol 38: 1329–1334.

28. Mathers AJ, Poulter M, Dirks D, Carroll J, Sifri CD, Hazen KC. 2014. Clinical Microbiology Costs for Methods of Active Surveillance for Klebsiella pneumoniae Carbapenemase-Producing Enterobacteriaceae. Infect Control Hosp Epidemiol 35:350–5.

29. Mathers AJ, Stoesser N, Sheppard AE, Pankhurst L, Giess A, Yeh AJ, Didelot X, Turner SD, Sebra R, Kasarskis A, Peto T, Crook D, Sifri CD. 2015. Klebsiella pneumoniae carbapenemase (KPC)-producing K. pneumoniae at a single institution: insights into endemicity from whole-genome sequencing. Antimicrob Agents Chemother 59:1656–63.

30. Wood DE, Salzberg SL. 2014. Kraken: ultrafast metagenomic sequence classification using exact alignments. Genome Biol 15:R46.

31. Ondov BD, Treangen TJ, Melsted P, Mallonee AB, Bergman NH, Koren S, Phillippy AM. 2016. Mash: fast genome and metagenome distance estimation using MinHash. Genome Biol 17:132.

32. Stoesser N, Sheppard AE, Peirano G, Anson LW, Pankhurst L, Sebra R, Phan HTT, Kasarskis A, Mathers AJ, Peto TEA, Bradford P, Motyl MR, Walker AS, Crook DW, Pitout JD. 2017. Genomic epidemiology of global Klebsiella pneumoniae carbapenemase (KPC)-producing Escherichia coli. Sci Rep 7:5917.

33. Nguyen LT, Schmidt HA, von Haeseler A, Minh BQ. 2015. IQ-TREE: a fast and effective stochastic algorithm for estimating maximum-likelihood phylogenies. Mol Biol Evol 32:268–74.

34. Sheppard AE SN, German-Mesner I, Vegesana K, Walker AS, Crook DW. 2018. TETyper: a bioinformatic pipeline for classifying variation and genetic contexts of transposable elements from short-read whole-genome sequencing data, vol 288001. Cold Spring Harbor, bioRxiv (Accecpted in Microbial Genomics).

